# Unbiased estimate of synonymous and non-synonymous substitution rates with non-stationary base composition

**DOI:** 10.1101/124925

**Authors:** Laurent Guéguen, Laurent Duret

## Abstract

The measure of synonymous and non-synonymous substitution rates (*dS* and *dN*) is useful for assessing selection operating on protein sequences or for investigating mutational processes affecting genomes. In particular, the ratio 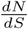 is expected to be a good proxy of *ω*, the probability of fixation of non-synonymous mutations relative to that of neutral mutations. Standard methods for estimating *dN, dS* or *ω* rely on the assumption that the base composition of sequences is at the equilibrium of the evolutionary process. In many clades, this assumption of stationarity is in fact incorrect, and we show here through simulations and through analyses of empirical data that non-stationarity biases the estimate of *dN, dS* and *ω*. We show that the bias in the estimate of *ω* can be fixed by explicitly considering non-stationarity in the modeling of codon evolution, in a maximum likelihood framework. Moreover, we propose an exact method of estimate of *dN* and *dS* on branches, based on stochastic mapping, that can take into account non-stationarity. This method can be directly applied to any kind of model of evolution of codons, as long as neutrality is clearly parameterized.

## 1 Introduction

The intensity and direction of selection operating on protein sequences can be evaluated by comparing the probability of fixation of non-synonymous mutations to that of neutral mutations. The ratio of fixation probabilities of non-synonymous vs. neutral mutations (denoted *ω*) is commonly estimated by comparing non-synonymous versus synonymous substitutions rates (denoted respectively *dN* and *dS*): under the assumption that selection on synonymous sites is negligible, the ratio 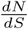 is expected to be a proxy for *ω*, and therefore to be informative on selective regimes on protein-coding sequences. Furthermore, the estimate of synonymous substitutions rates can also be useful in itself, e.g. to be used as a molecular clock, or to investigate variation in mutation rates or biased-gene conversion along genomes.

Substitution rates (*dN* and *dS*) are expressed in terms of number of (non-)synonymous substitutions per (non-)synonymous site. One important issue is therefore to quantify the number of (non-)synonymous sites. Historically, the first methods developed to estimate *dS* and *dN*, directly compared sequences to count the numbers of synonymous and non-synonymous substitutions, and used elaborate formula to account for the “per (non-)substitution site” feature (Li *et al*., 1985; Nei and Gojobori, 1986).

Subsequent methods relied on sequence alignments in a phylogenetic context, and the maximum likelihood of probabilistic codon-based substitution models on these alignments (Goldman and Yang, 1994; Yang and Nielsen, 2000; Guindon *et al*., 2004; Kosakovsky Pond *et al*., 2005; Yang, 2007). Once *ω* has been estimated by maximum likelihood, *dN* and *dS* can also be inferred ancestral sequence reconstruction: on each branch, the number of (non-)synonymous substitutions is estimated, and to consider the “per (non-)synonymous site” feature, the expected numbers of (non)-synomynous neutral substitutions are estimated by applying a similar but neutral model, *i.e*. without selection (Goldman and Yang, 1994; Yang and Nielsen, 2000; Kosakovsky Pond and Frost, 2005). In addition to the estimate of *dN* and *dS*, such an approach is a convenient way to access branch-specific and site-specific substitution process, specifically while looking for signals of episodic, positive selection (Messier and Stewart, 1997; Kosakovsky Pond and Frost, 2005; Lemey *et al*., 2012).

But up to now, programs used to compute *dN* and *dS* have two drawbacks. First, they propose approximate computations of the counting of the numbers needed, for the counting of effective (non-)synonymous substitutions as well as for the normalization “per (non-)synonymous site”. For example, in (Kosakovsky Pond and Frost, 2005), Kosakovsky-Pond and Frost consider the most parsimonious substitution scenarios between expected ancestral states at top and bottom of the branches, and compute which part of each scenario is synonymous or not. Afterwards, they use an inferred model and its neutral equivalent to estimate *dN* and *dS*. However, choosing a given substitution scenario (the most parsimonious, or even the most likely one) will forget many other possible scenarios, especially when the branch gets longer and the selection gets smaller.

Second, these programs assume stationarity in the modeling of the data, i.e. assume that codon frequencies are constant all along the evolutionary process. It is now well established that in many cases this assumption is false. For example, in mammals, genomic landscapes are characterized by large-scale variation in GC-content along chromosomes (the so-called isochores) (Bernardi *et al*., 1985), which are caused by the process of GC-biased gene conversion (gBGC) (Duret and Galtier, 2009). Variation in the intensity of gBGC among taxa (notably due to variation in recombination rate) caused frequent changes in gene GC-content along the mammalian phylogeny (Romiguier *et al*., 2010). Variations in GC-content are also frequently observed in bacteria, notably during the reductive genome evolution of endosymbionts such as *Buchnera aphidicola* (van Ham *et al*., 2003; Moran *et al*., 2008; Moran, 1996; Pérez-Brocal *et al*., 2006), but also in free-living organisms such as *Prochlorococcus marinus* (Rocap *et al*., 2003; Dufresne *et al*., 2005; Yu *et al*., 2012; Dufresne *et al*., 2003; Paul *et al*., 2010). These changes in GC-content affect both codon (Wernegreen and Moran, 1999; Moran, 1996) and amino-acid (Mouchiroud *et al*., 1991; Wernegreen and Moran, 1999; Itoh *et al*., 2002; Moran *et al*., 2008) frequencies.

In this article, we illustrate through simulations how assuming stationarity leads to a systematic bias in *dN, dS* and 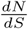 estimates, and we show that this bias can be properly removed when stationarity assumption is released. Next, we introduce a new method based on stochastic mapping for an accurate estimate of *dN* and *dS*. Instead of choosing a given scenario between pairs of ancestral states on branches, this method integrates over all possible scenarios, in accordance with their probability given the model and the length of the branch, to compute more precisely *dN* and *dS*, following the definition given in Kosakovsky Pond and Frost (2005). We implemented this method in bio++ libraries (Guéguen *et al*., 2013), so that it can be used without any constraint of stationarity or homogeneity of the process, and can give access to branch and/or site specific estimates. Using this method, we explore the bias induced by the assumption of stationarity on the estimates of *dN, dS* and 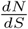, and show that this bias is fixed with our method. Finally, an application of this method and the importance of considering non-stationarity are illustrated on a set of orthologous primate genes.

## 2 Stochastic Mapping

Stochastic mapping is a way to infer substitution events based on probabilistic modeling estimates. In 2002, Rasmus Nielsen proposed a bayesian approach to map substitution events on the branches of a phylogenetic tree, given a probabilistic substitution model (Nielsen, 2002). Since then, many theoretical and computational works have been made to describe accurately the substitution process along a phylogenetic tree, given a probabilistic model and a sequence alignment (Ball and Milne, 2005; Dutheil *et al*., 2005; Minin and Suchard, 2008; Hobolth and Stone, 2009).

These works are based on computing the expected number of substitution events of a given category along a branch. These estimates are conditioned by the states at both ends of this branch. Moreover, Minin and Suchard have proposed a way to compute the expected time spent in a given state on this branch, under the same conditions (Minin and Suchard, 2008). With real data, the sequences on the ancestral nodes are not known, but it is possible to compute the posterior expectations on each branch given the data and the substitution process (Romiguier *et al*., 2012).

Hereafter, we use this methodology specifically on two categories of events: synonymous and non-synonymous substitutions. Similarly to the “per (non-)synonymous site” normalization for *dN* and *dS*, the expected numbers of substitution events on a branch have to be corrected given the changing ancestral sequence all along the branch. (O’Brien *et al*., 2009) where 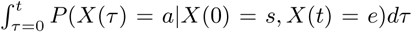 is the time spent by *X* in state *a* during duration *t*, given the initial state *X (0)* = *s* and the final state *X* (*t*) = *e*. We can define the conditional **ability** of process *X* based on model ℳ, under the same conditions, to perform ℒ—events:

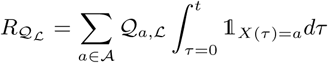

given that *X*(0) = *s, X*(*t*) = *e*. In Minin and Suchard (2008), this value is defined as the reward of vector 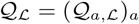, which conditional expectation 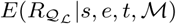 can be computed in the same way as 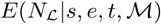.

For any branch *b* ∈ *T* of length *t*_b_ and any site *i*, we can compute *P*(*s,e*|*b,D*_*i*_, ℳ) the probability that the state of site *i* at the top (resp. bottom) of *b* is *s* (resp. *e*) given ℳ and data *D*_i_ on this site. Then the expected *a posteriori* number of ℒ—events on site *i* in branch *b* is *E*(*N*_ℒ_|*b, D*_i_, ℳ) *= Σ*_s e_ *E*(*N*_ℒ_|*s, e, t*_*b*_).*P*(*s, e*|*b, D*_i_, ℳ), which sums up on the sites: *E*(*N*_ℒ_|*b,D*, ℳ) = Σ_i_ *E*(*N*_ℒ_|*b,D*_i_, ℳ) to obtain the *a posteriori* expected count of ℒ—events on branch *b* given the data *D* and the model ℳ.

In a same manner, we can compute the expected ability of any model ℳ′ (with generator *Q*′) to perform ℒ—events given the *a posteriori* probabilities of sequences given *D* and *M*: 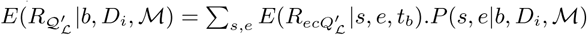 and 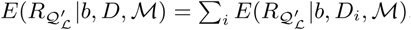.

Here these expectations are computed on all scenarios given the data and model ℳ, which is in theory the true model, and in practice will be the most likely model.

How can we use this definition to compute relevant *dN* and *dS*? At each time *τ*, the property of a site to be (non-)synonymous is based on the rates of all the (non-)synonymous substitutions this site can undergo. These rates depend on the considered model. For example, a site with the codon `AAA` (which only synonymous codon is `AAG`) will be more synonymous with a model that favors `A` and `G` nucleotides than with a model that favors `C` and `T`. So these rates will have to be computed with a model similar to ℳ, but defined as neutral, i.e. which does not favor synonymous or non-synonymous substitutions in its definition (Yang and Nielsen, 2000; Kosakovsky Pond and Frost, 2005). Hence, to compute relevant *dN* and *dS*, the expected counts of (non-)synonymous substitutions will be normalized considering the number of potential substitutions expected according to the same model, but without selection. If ℳ is the model used to describe the process, with generator 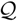, we denote ℳ^0^ its neutral version, with generator 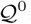. In the case of model YN98 (Yang and Nielsen, 1998), 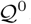 is the same as 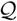, with *ω* = 1. And the expected number of (non)-synonymous substitutions that would have been performed by ℳ_0_ is by definition the ability of model ℳ_0_ along the history defined by *D* and ℳ.

The ratio 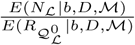 is then considered as an *a posteriori* normalized count of the ℒ—events on branch b. Since the models are built on codon sequences, they are normalized such that there is one substitution per codon per unit of time on sequences at equilibrium. It is then straightforward to see that the ability of a model to perform any substitution equals 1 per unit of time per codon. In the case of *dN* and *dS*, the normalization is not “per codon” but “per nucleotide”, which means the ability of a model to perform any substitution should be 1 per unit of time per nucleotide, i.e. 3 times the previous one. Finally, we obtain the equivalents of *dN* and *dS* in the methodology of stochastic mapping: 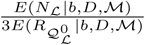.

## 3 Applications

To investigate the bias induced by stationarity assumption, we use stochastic mapping to compute relevant *dN, dS* and 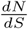 estimates on simulated and empirical sequence datasets, both of which are subject to changes in GC content. The same model has been used, aka the model proposed in Yang and Nielsen (1998) (denoted “YN98”), both in homogeneous and non-homogeneous (or branch) modelings. To model a non-stationary process, it is necessary to introduce root codon frequencies. To reduce the number of parameters to estimate, root and equilibrium codon frequencies are computed as products of position nucleotide frequencies instead of a full parametrization of the codon frequencies (61 parameters). In simulations, nucleotide frequencies are considered as identical for all positions (denoted “F1X4”, with 2 × 3 parameters). For real dataset analyses, nucleotide frequencies are position specific inside codons (denoted “F3X4”, with 2 × 9 parameters, because of 3 equilibrium frequencies), and normalized as stop codon frequencies are set to 0.

In a first step, parameter *ω* is estimated through maximum likelihood computation of model, root frequencies and branch lengths on each alignment. Then, in a second step, *dN* and *dS* are computed using normalized stochastic mapping as described in Method section, from this optimized model and tree.

This procedure has been implemented in the Bio++ program suite (Guéguen *et al*., 2013). It can then easily be used on the numerous models that are available in this suite, and most importantly in any non-homogeneous modeling. Moreover, it can output both site-specific and/or branch-specific estimates.

This suite was used for simulations, maximum likelihood estimates and stochastic mapping computations. We also performed the same estimates under stationary assumption with codeml (Yang and Nielsen, 2000), and the results exhibit similar biases to those obtained with our approach (see Fig. S1 in supplementary material).

## 4 Data

### 4.1 Simulated dataset

To study the influence of the non-stationarity in G+C content on the maximum likelihood estimate of *ω*, we simulated the evolution of 100 coding sequences of 3000 codons. Each simulation started from an ancestral sequence with determined proportion of G+C, noted *θ*_root_, and ran along the tree depicted in Figure 1, using an homogeneous YN98+F1X4 model with determined G+C equilibrium frequency, noted *θ*_*eq*_. Each θ value (*θ*_eq_ and *θ*_root_) ranged from 0.1 to 0.9 per step of 0.1. We simulated negative, weakly negative, neutral, and weakly positive selection (resp. *ω* = 0.1, *ω* = 0.9, *ω* = 1, *ω* = 1.1).

**Figure 1:**
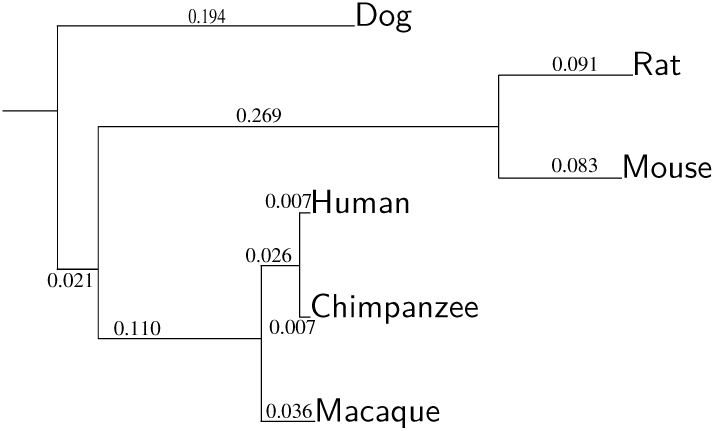
Phylogeny of the studied species in the mammalian dataset. The same tree is used for simulation with two different theta values: *θ*_root_ is the G+C probability at the root, *θ*_*eq*_ is the equilibrium G+C probability.

### 4.2 Mammalian dataset

From the data studied in Kosiol *et al*. (2008), we retrieved 6055 sequence alignments of orthologous genes present in human, chimpanzee, macaque, mouse, rat and dog genomes.

## 5 Results

### 5.1 Assessment on simulated data

On all data, we inferred the most likely model, which gave the estimate of *ω*, and then we used our approach to estimate *dN, dS* and 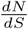.

When *ω* is estimated with a stationary model, decreasing G+C content along the tree entails a systematic over-estimate of *ω*, and increasing G+C content entails an systematic under-estimate of *ω*, whereas the same estimate without the hypothesis of stationarity are not biased (see Figure 2, results obtained with other *ω* are shown in supplementary figures S2 to S4). These under or over estimates can lead to false qualitative interpretation of selection, as dubious positive selection can be inferred in case of decreasing GC-content, or dubious negative selection in case of increasing GC-content (as illustrated in simulations with neutral and nearly-neutral models, see supplementary figures S5 to S10).

**Figure 2:**
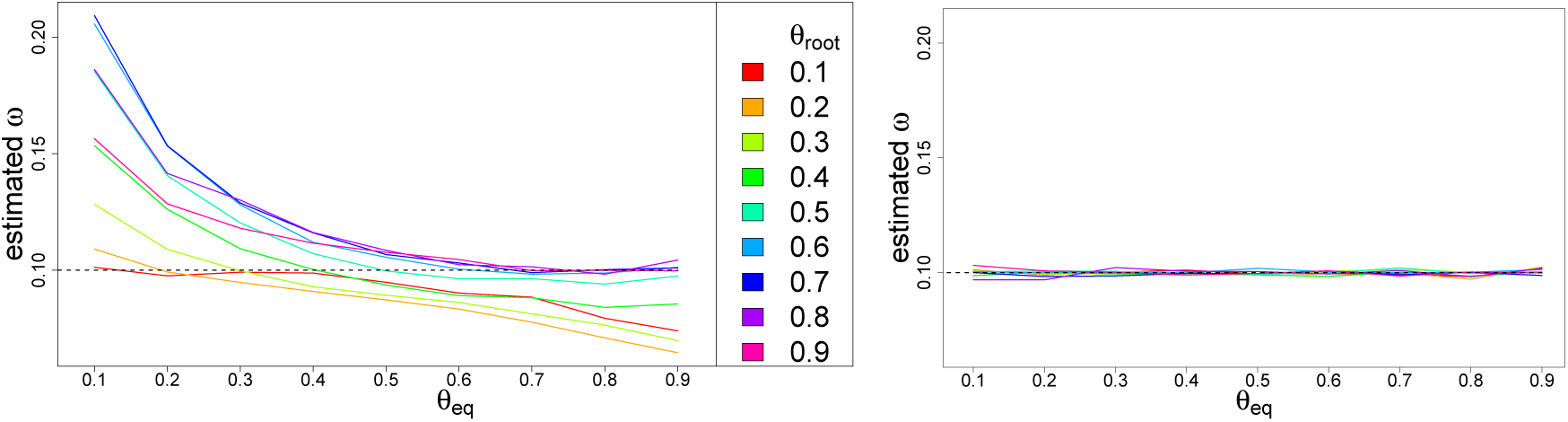
**Estimate of** *ω* = 0.1 with a stationary model (left) and non-stationary model (right), on simulated data with changing G+C content. *θ*_root_: G+C frequency in the root sequence. *θ*_eq_: G+C equilibrium frequency of the simulation model.

For the estimate of dN and dS, assuming stationarity biases both the estimates of *dN* and *dS* in similar ways (Figure 3). These values are mostly under-estimated (Fig. 4) when equilibrium GC is very different from 0.5 and GC content changes (either up or down). This means that in these cases the inferred trees are too short. We also observe that unbiased estimates of *dN* and *dS* decrease with equilibrium GC content. But this is not due to our method, since on stationary processes estimates of *dN* and *dS* computed with codeml have a similar trend (see the dashed line in Fig. S1 in supplementary material).

**Figure 3:**
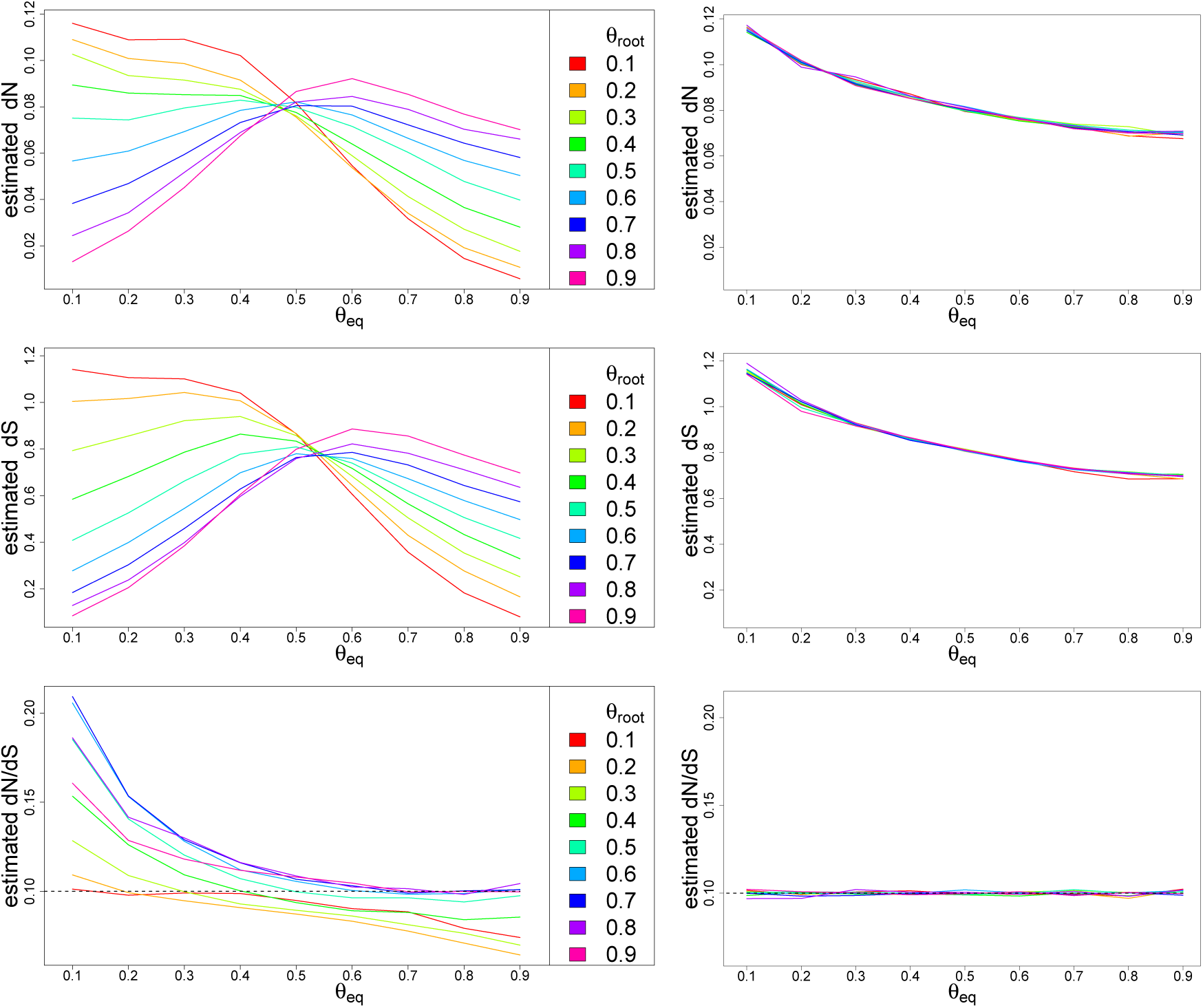
**Estimate of *dN, dS* and** 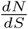 with a stationary model (left) and non-stationary model (right), on simulated data with changing G+C content and *ω* = 0:1. *θ*_root_: G+C frequency in the root sequence. *θ*_eq_: G+C equilibrium frequency of the simulation model.

**Figure 4:**
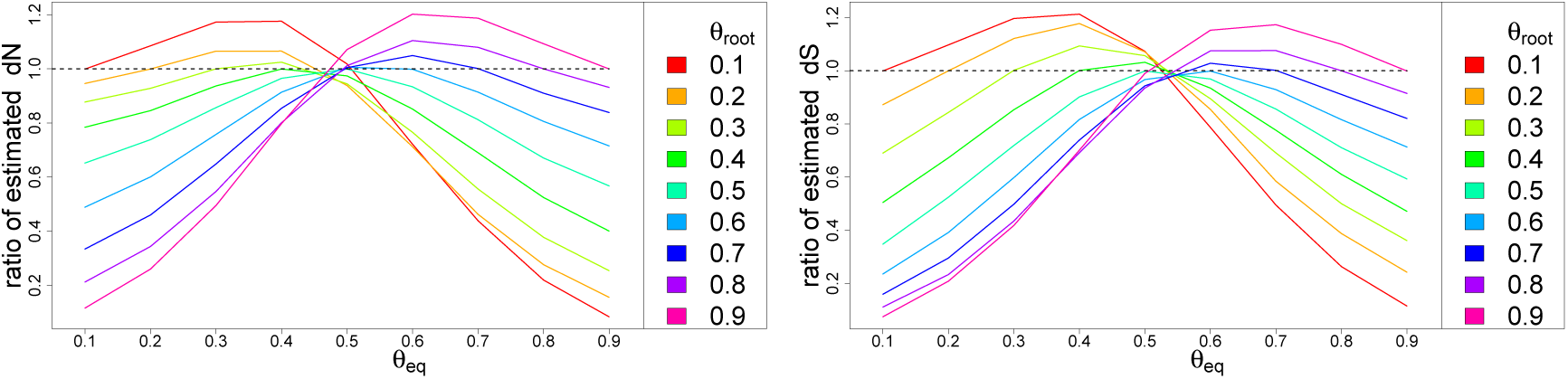
Ratio of substitution rates estimated with stationary model over substitution rates estimated with non-stationary model. Sequences were simulated with changing G+C content and *ω* = 0.1. Left: dN. Right: *dS*. *θ*_root_: G+C frequency in the root sequence. *θ*_eq_: G+C equilibrium frequency of the simulation model.

Actually, when the dynamics of GC content is heterogeneous, the bias is not systematically in the same direction whether GC increases (or decreases), but it depends also on the GC of other branches, since a stationary modeling (hence homogeneous) will estimate its GC equilibrium from all branches. For example, on the same tree, we considered a model with stationary GC from the root to the primate leaves, and changing GC on the branches leading to dog and to rodents. As shown in Figure 5, estimates of 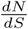 on primate branches are biased with the hypothesis of stationarity, even though the process is indeed stationary on these branches. But the non-stationarity on the other branches misleads the estimated stationary model.

**Figure 5:**
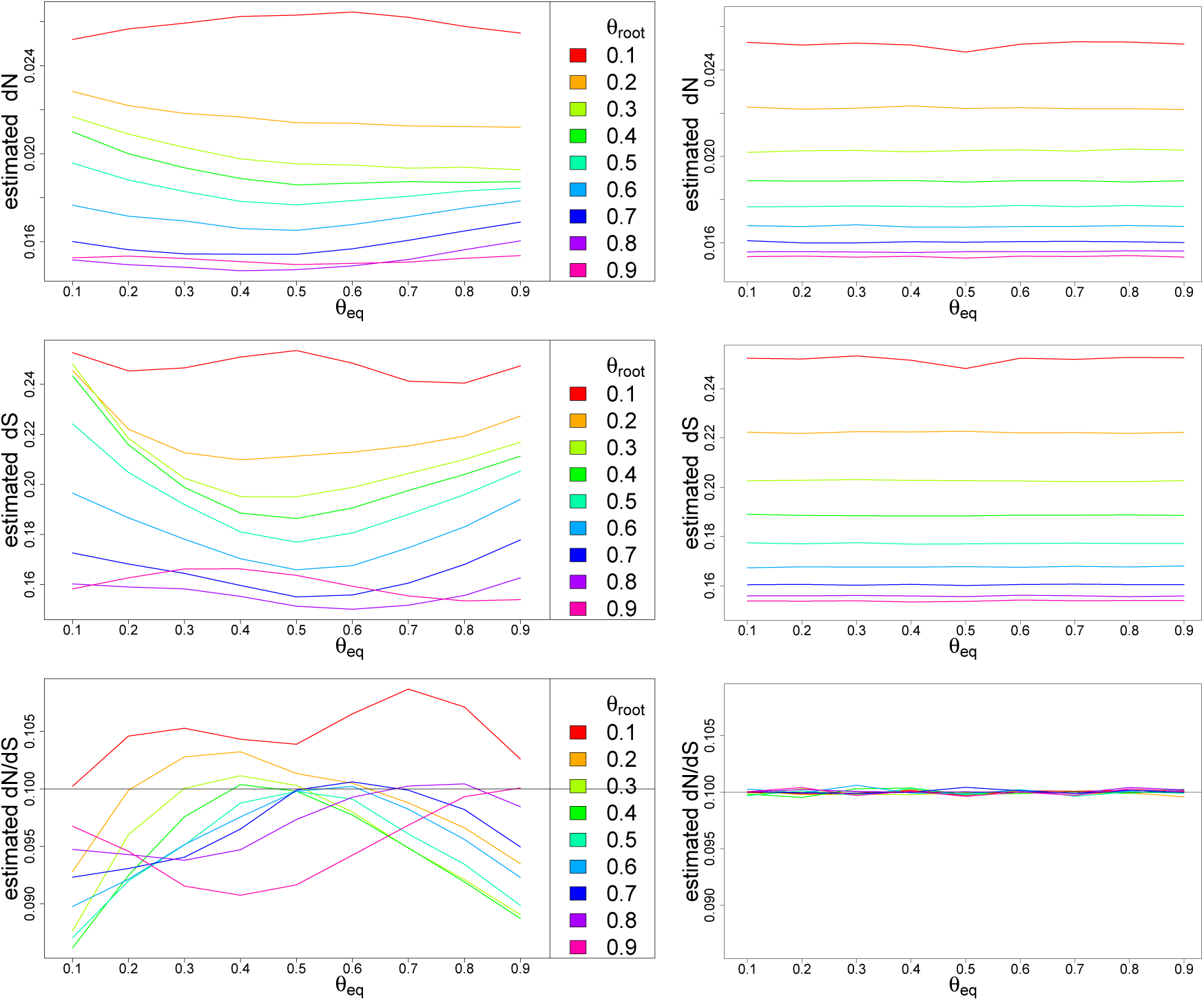
**Estimate of** *dN, dS* **and** 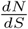 **on primate branches** with a stationary model (left), and a non-stationary non-homogeneous model (right), on simulated data with changing G+C content on dog and rodent branches, and *ω* = 0:1. *θ*_root_: G+C frequency in the root and primates sequences. *θ*_eq_: G+C equilibrium frequency of the simulation model on dog and rodent branches.

### 5.2 Study on mammalian data set

We performed two different maximum likelihood estimates of the mammalian data set: a stationary homogeneous YN98+F3X4 model (21 branch and model parameters), and a non-stationary non-homogeneous model (31 additional parameters) with three homogeneous YN98 models, one for the primate clade, one for the rodent clade and one for the dog branch. We used three models to match the heterogeneity in equilibrium GC content found between these clades Romiguier *et al*. (2010). We computed *dN* (resp. *dS*) in the primate clade by summing the stochastic mapping *dN* (resp. *dS*) of all branches of this clade.

Since the modelings are nested, we performed likelihood ratio tests on all estimates, and corrected multiple testing using Benjamini-Hochberg correction. The increase in likelihood is significant (using an LRT test with 31 degrees of freedom) with an 1% FDR value, in 83.4% of the genes (Fig. 6).

**Figure 6:**
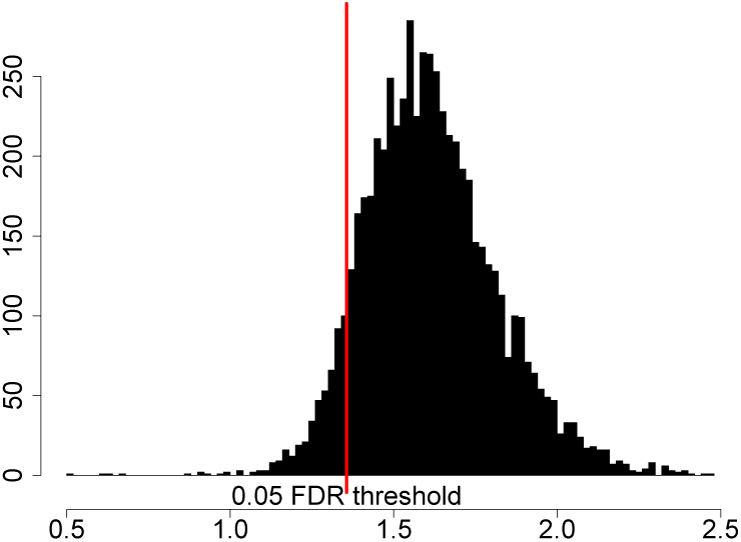
Log10 of the differences in log-likelihoods between stationary and non stationary models on mammalian data. The red line stands for the 5% FDR threshold.

If we compare the estimates of stationary versus non-stationary modeling, we see that the estimates of *dN* are mostly lower, but not correlated with the evolution of GC-content at third codon position (GC3) (Figure 7). On the contrary, we see an influence of the evolution in GC3 on the bias in the estimate of *dS*, and then a more important under-estimate of 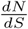 with genes far from stationarity in GC3. As noticed in the simulation section, the bias is not correlated with the sign of change in GC3 because we performed a non-homogeneous modeling, and the bias depends also of the evolution of GC content in the other branches. However, the effect is quite noticeable, the relative error on *ω* estimate is at least 10% for 59% of the genes, or at least 33% for 13.4% of the genes (Figure 8).

**Figure 7:**
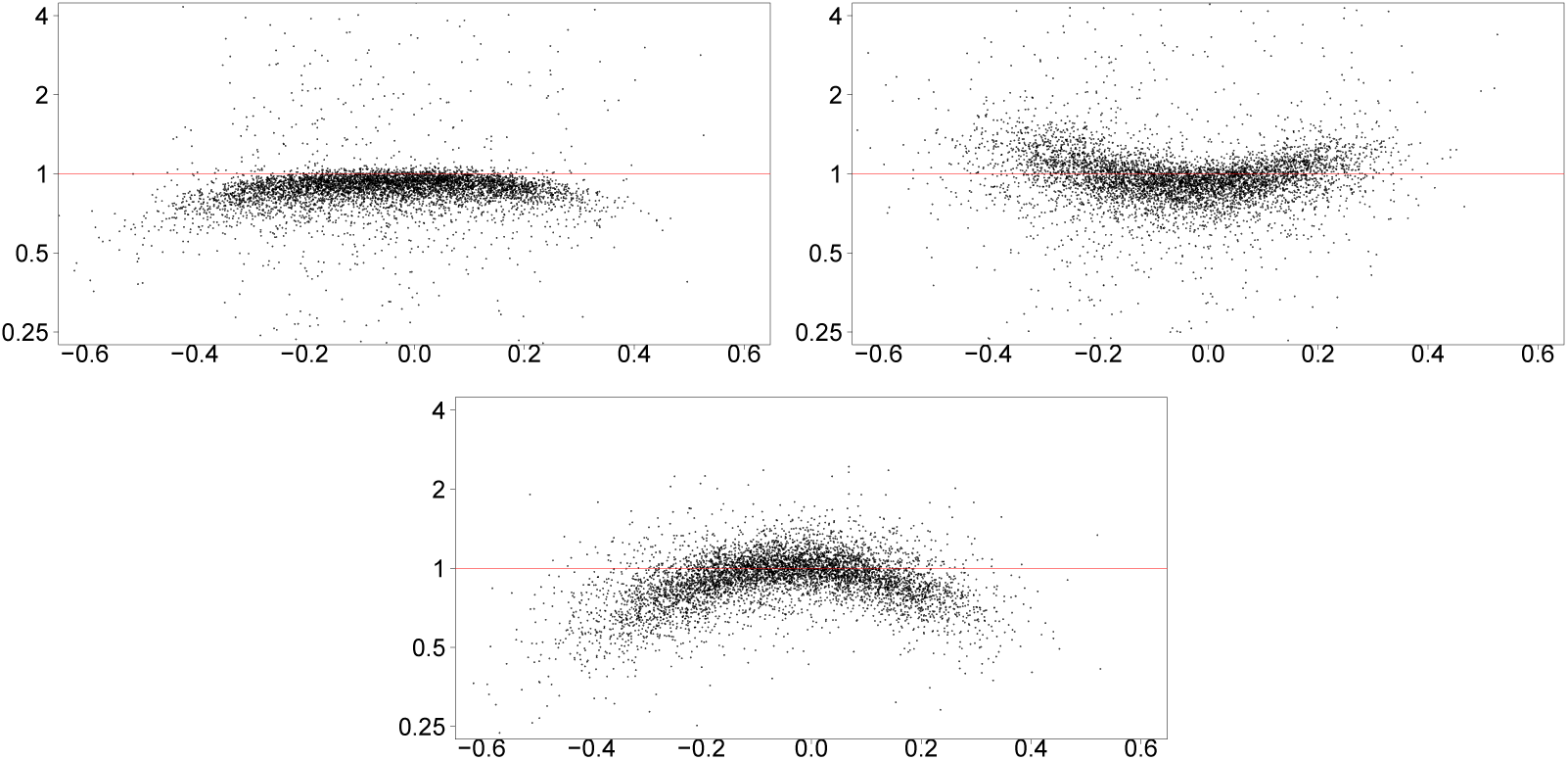
**log2 of the ratios of estimates of *dN, dS* and** *dN* = *dS* with a stationary model over the estimates with a non-stationary model, according to the change in *GC*3 content in the primate clade.

**Figure 8:**
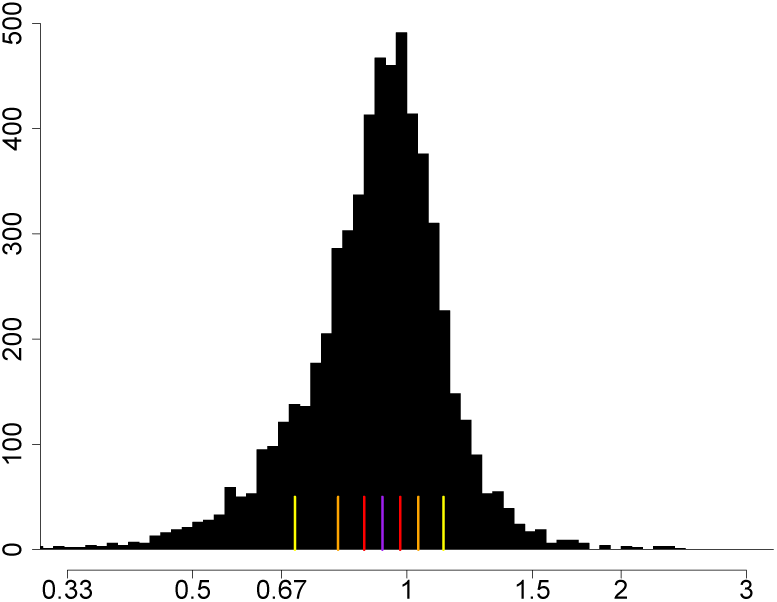
**Histogram of the ratios** in estimates of *ω* in stationary model over non-stationary model on mammalian data. Yellow, orange and red lines stand for 12:5%, 25% and 37:5% quantiles. The purple line represents the median.

### 6 Conclusion

Our analyses, both on simulated and empirical datasets, show that estimates of *dN, dS* and *ω* can be biased when using standard methods, which assume sequence stationarity. The strength of the bias depends on the gap between the equilibrium and the actual base composition. Generally, estimates of *ω* are more robust to this bias than those of *dN* or *dS* (Fig. 3), but in extreme cases, our simulations showed a two-fold difference between the true and estimated value of *ω*. This bias can have a profound impact for analyses aiming at comparing average values of *ω* among large gene sets. For instance, to investigate the parameters that explain variations in the efficacy of selection, many studies have compared the genome-wide average of *ω* across different taxa (e.g. Galtier (2016)). The genome-wide average of *ω* varies from 0.13 to 0.17 among 48 bird species (Weber *et al*., 2014), and from 0.10 to 0.22 among 106 amniote species (Figuet *et al*., 2016). Thus, at this scale, systematic errors in the estimate of *ω* caused by differences in the equilibrium base composition along lineages might have an important impact on observed patterns.

The method that we developed, based on stochastic substitution mapping, provides unbiased estimates of *dN, dS* and 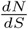. Moreover, this method can be used with any type of codon modeling, as long as it is possible to define a neutral model with given parameter values.

Substitution mapping counts events on each branch and on each site. We used it in the estimate of *ω* on the whole tree or on a set of branches, but it can be used on any specific branch, to look for episodic selection regimes. It seems also straightforward to adapt this approach in the estimate of selection on specific sites, and indeed it is already possible with “simple” models such as the ones we considered in this article. However, most site-specific studies consider site-models to model the heterogeneity in selection along the sequence (Yang *et al*., 2000). Results of substitution mapping depend on the model used, and it seems reasonable to use similar site-models in the case of heterogeneous sequences.

As described in (Minin and Suchard, 2008), in addition of computing the expectation of counts and times on branches, it is possible to compute their variance (and other moments). This would provide statistical information on the accuracy of the estimates of *dN* and *d*S, and we expect that it will be quite important in site specific estimates, since site specific data is quite poor, and far more in restrained clades.

### 7 Availability

Our method has been implemented in the Bio++ suite (Guéguen *et al*., 2013). The maximum likelihood program is called bppml, and is available at the address http://biopp.univ-montp2.fr/BppSuite. The stochastic mapping program is called mapnh, and is available at the address http://biopp.univ-montp2.fr/forge/testnh.

A short tutorial about model inference and stochastic mapping as described in this article is available there: https://github.com/BioPP/supp-mat/tree/selection_vs_GC.

## 8 Acknowledgements

We thank Marie Sémon for her comments on the manuscript. This work was supported by the Centre National de la Recherche Scientifique, and by the Agence Nationale de la Recherche (Ancestrome: ANR-10-BINF-01-01). This work was performed using the computing facilities of the CC LBBE/PRABI.

